# A Bayesian approach for correcting Tn5 transposition bias in ATAC-seq footprinting

**DOI:** 10.1101/525808

**Authors:** Vinayak V. Viswanadham, Shiv S. Pillai, Vinay S. Mahajan

## Abstract

ATAC-seq exploits the observation that the pattern of transposition of a hyperactive Tn5 transposase in native chromatin mirrors genome-wide chromatin accessibility. It has been suggested that transposition observed around transcription factor binding motifs can be used to assess their occupancy in the form of footprints. However, we show that the vast majority of footprints observed at transcription factor motifs in ATAC-seq data spuriously arise from the intrinsic sequence-dependent transposition site bias of Tn5 and are also observed in naked DNA. We demonstrate that the Tn5 transposition bias can be corrected using existing tools for sequence bias correction and a novel estimate of global occupancy in order to produce more reliable estimates of footprints.

---

ATAC-seq has emerged as a powerful method to assess genome-wide DNA accessibility in native chromatin ^1^. This approach relies upon the use of a hyperactive variant of the Tn5 transposase that can cleave and tag DNA for next-generation sequencing in a single step dubbed tagmentation. Tn5 transposition in native chromatin is largely restricted to accessible regions, resulting in a rapid method to map genome-wide chromatin accessibility. Footprints of DNA-bound transcription factors (TFs) that occlude Tn5 transposition can be observed in ATAC-seq data ^1^. Such footprints serve as a surrogate measure of transcription factor occupancy and this approach is conceptually identical to DNAse I footprinting ^2^.

In recent studies, bioinformatic tools such as CENTIPEDE have been applied to generate a probabilistic estimate of footprinting at individual instances of a TF motif in ATAC-seq data ^3^. We used CENTIPEDE to examine transposition footprints at matches to all the JASPAR core non-redundant vertebrate motifs found in the ATAC-Seq dataset generated from a B cell line initially described in Buenrostro et al ^1,4^ We found that several motifs were estimated to be footprinted by CENTIPEDE although the transposition pattern surrounding these motifs showed no obvious indication of footprinting (Figure 1). This is not surprising given that CENTIPEDE does not explicitly model the occurrence of diminished transposition at motif sites. Instead, the CENTIPEDE model distinguishes genomic intervals that have a *non-uniform* distribution of insertion sites within the interval with a high coverage relative to a background of *uniformly* transposed intervals with lower coverage. We illustrate this using simulated data (Figure 2), where transposition events are modeled from a normal distribution centered on a motif, such that there is no discernible TF footprint. We find that as might be expected from the model specifications, CENTIPEDE functions primarily as a coverage detector. This also explains the correlation between CENTIPEDE footprinting predictions and ChIP-Seq data. TF binding and unbinding occurs on a much smaller timescale than the consequent increase in chromatin accessibility, which is sustained by the recruitment of numerous chromatin modifying enzymes ^5^. While ChIP-seq detects TF binding, ATAC-Seq identifies consequent changes in accessibility and the degree of chromatin accessibility is proportional to coverage in ATAC-Seq tracks. However, as previously seen with DNAse-Seq, TFs which exhibit a high motif occupancy rate due to a high affinity for target DNA are more likely to be footprinted in the conventional sense and the depth of their footprint in ATAC-Seq data can be expected to be proportional to their residence time ^6^. CTCF falls into the category of TFs with relatively long residence times, and we explored if other similar TFs can be identified by examining transposition patterns at TF motif matches in ATAC-Seq data. Due to the limitations of the CENTIPEDE algorithm described above, we defined a footprint score that is explicitly based upon the magnitude of reduction in the number of cut sites within the motif matches with respect to the adjacent flanking intervals (Figure 3).

**FIGURE 1:**
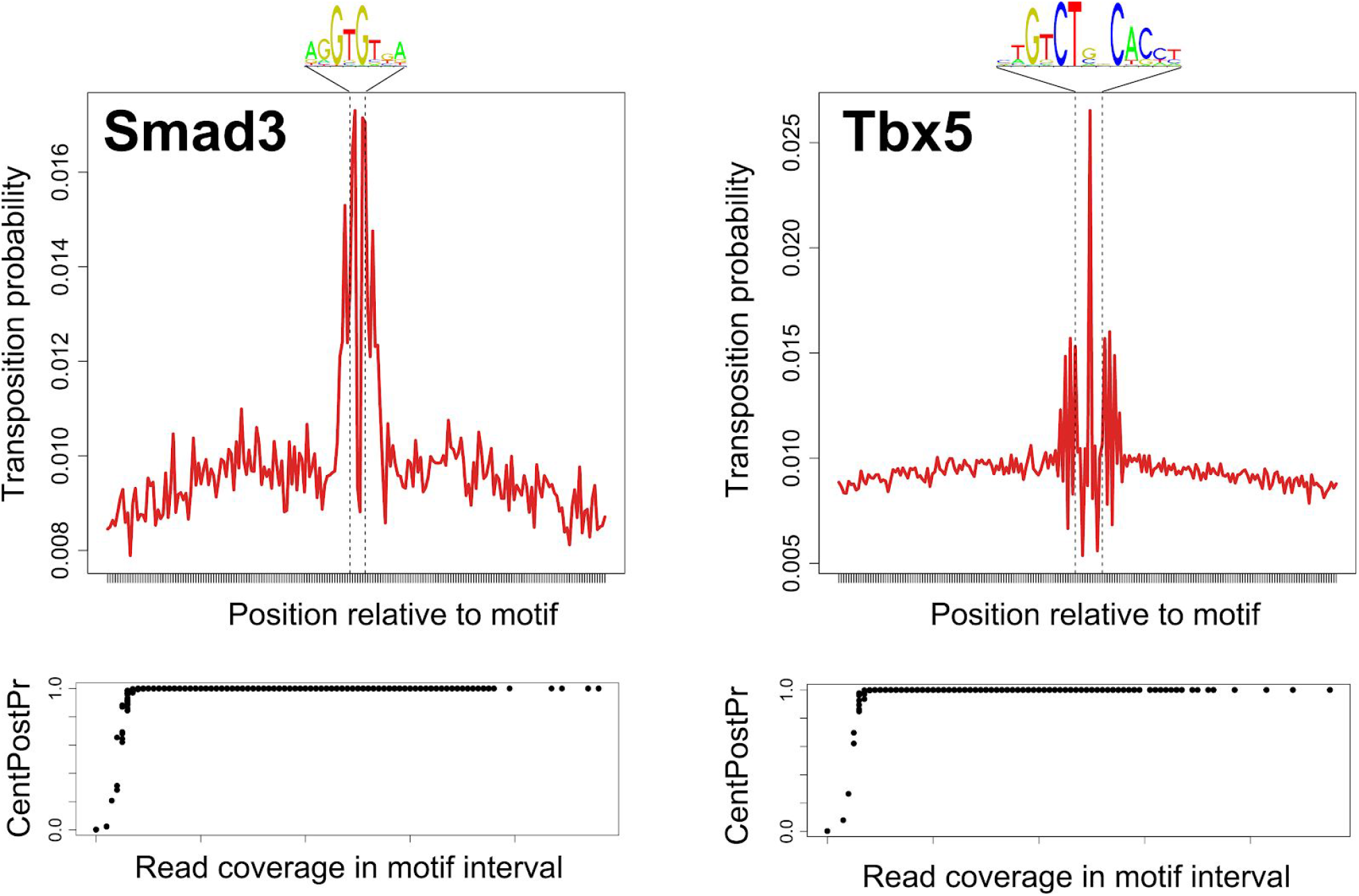
Transposition patterns surrounding motifs in ATAC-Seq accessible intervals deemed footprinted by CENTIPEDE. Transposition pattern around all instances of Smad3 (2383 matches) and Tbx5 (6830 matches) motif matches in ATAC-Seq accessible intervals (+/− 100 bp were considered). Transposition probability is calculated as the proportion of intervals carrying a transposition event at the given position relative to the motif match. The lower panel shows the relationship between CENTIPEDE posterior probability (CentPostPr) of footprinting and read coverage (i.e. number of transposition events) per motif-containing interval. Despite no visual indication of diminished transposition at the motif matches, over 90% of motif-containing intervals are deemed footprinted (CentPostPr = 1) by the CENTIPEDE algorithm.

**FIGURE 2:**
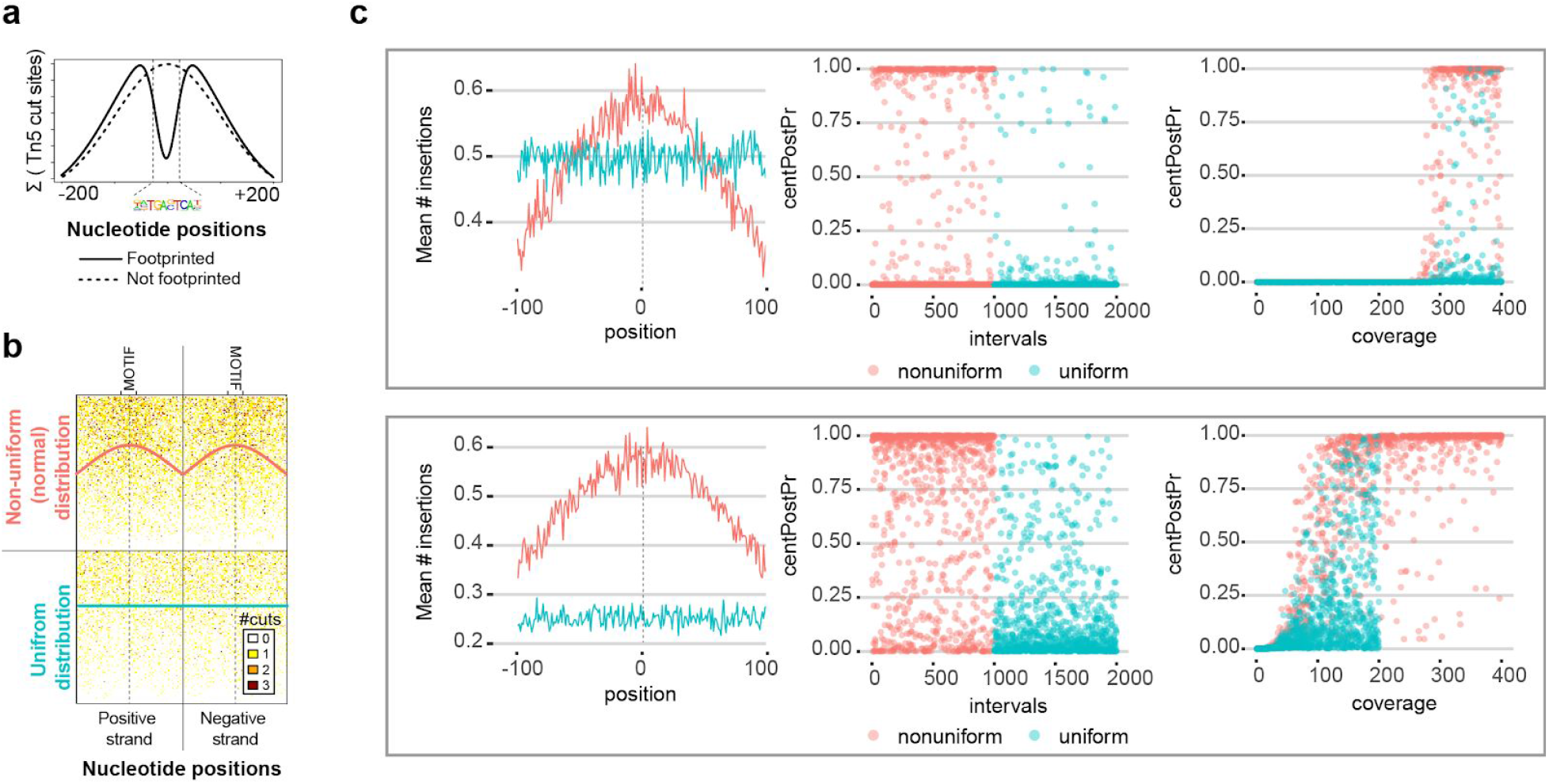
CENTIPEDE footprint estimates do not reflect steric occlusion at motif matches. **(a)** Schematic depiction of ATAC-seq transposition sites surrounding a motif that would be deemed footprinted (solid line) or not footprinted (dashed line). The positional distribution of cut sites surrounding a non-footprinted motif in ATAC-seq data can be approximated by a normal distribution centered on the motif. **(b)** Simulated data of cut site counts across 2000 intervals bearing a non-footprinted motif to test CENTIPEDE footprint prediction is shown. The cut sites in each interval were drawn from a normal distribution centered on the motif site (pink) in the top half, and from a uniform distribution (blue) in the lower half. Each row represents cut sites across an interval. The total cut site coverage for each interval was randomly sampled from a uniform distribution ranging from 0 to 400 cuts. Two sets of data were simulated: (i) with no difference and (ii) with a 2-fold difference in the mean coverage between the intervals bearing a uniform and a non-uniform positional distribution. Only the former is shown here. **(c)** CENTIPEDE predicts non-uniform intervals (pink) as having a greater posterior probability (centPostPr) of being footprinted than uniform intervals (blue), despite all intervals being explicitly modeled as not exhibiting any footprint. Moreover, higher probabilities are calculated for intervals with greater coverage. This effect is especially pronounced when the non-uniformly distributed intervals have a higher coverage. Two simulated data sets are shown: (i) with no difference (top panel) and (ii) with a 2-fold difference (lower panel) in the mean coverage between the intervals bearing a uniform and a non-uniform cut site distribution.

**FIGURE 3:**
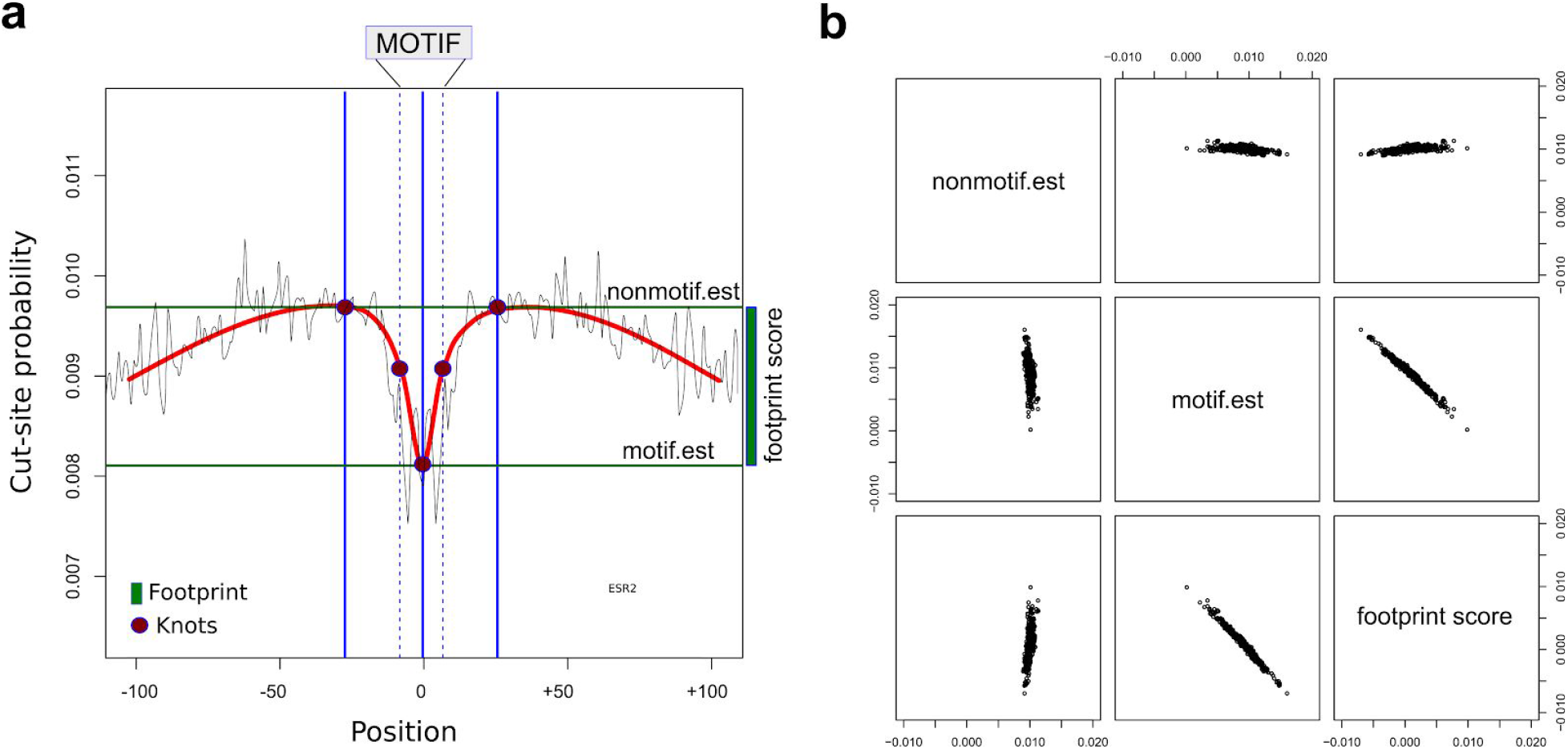
Calculation of footprint scores. **(a)** A spline function was fit to get a smoothed estimate of position-specific cut-site probabilities with knots placed at the midpoint of the motif, the ends of the motif match, and +/− 19 bases from the motif. The magnitude of the footprint was defined as the difference (green bar) in the cut-site probability at the knots placed in the flanking intervals and the center of the motif. **(b)** Footprint scores are not correlated with transposition frequencies at the spline knot positions placed at the 19-base flanks (“non-motif.est”), implying that the footprint scores are not correlated with read coverage.

Although Tn5 transposition is random enough for most biological applications, Tn5 is known to exhibit a small sequence bias in its target sequences ^1,7,8^. In order to estimate the transposition bias in Tn5-tagmented libraries, we calculated the consensus sequence around the read start sites in a whole-genome sequencing library prepared from naked human genomic DNA using Tn5 tagmentation (Amini et al. 2014) and in an ATAC-Seq library prepared from native chromatin ^1^. To minimize differences in the sequence space that was sampled for this comparison, we restricted our analysis only to the genomic intervals that were accessible in the ATAC-seq library ^1^. Both libraries revealed a subtle but highly conserved palindromic 19-base long nucleotide sequence bias around the read start sites (Figure 4a). Of note, no bias is observed when the insertion sites are randomly offset by up to 10bp, and the bias is present whether reads within different distances of ATAC-seq peak summits are considered (Figure S1), reads are filtered for MAPQ scores, or reads are downsampled (data not shown). The symmetry in the bias observed in the top and bottom strands of the target DNA is concordant with the homodimeric structure of the Tn5 transposase comprising two active sites and is likely to result from the residues that contact target DNA (Figure 4b). We speculate that the slight difference in the Tn5 transposition bias between naked DNA and native chromatin may arise from structural differences in naked genomic DNA and chromatin (e.g. supercoiling, effect of nucleosomes) or from possible differences in experimental conditions.

**Figure 4:**
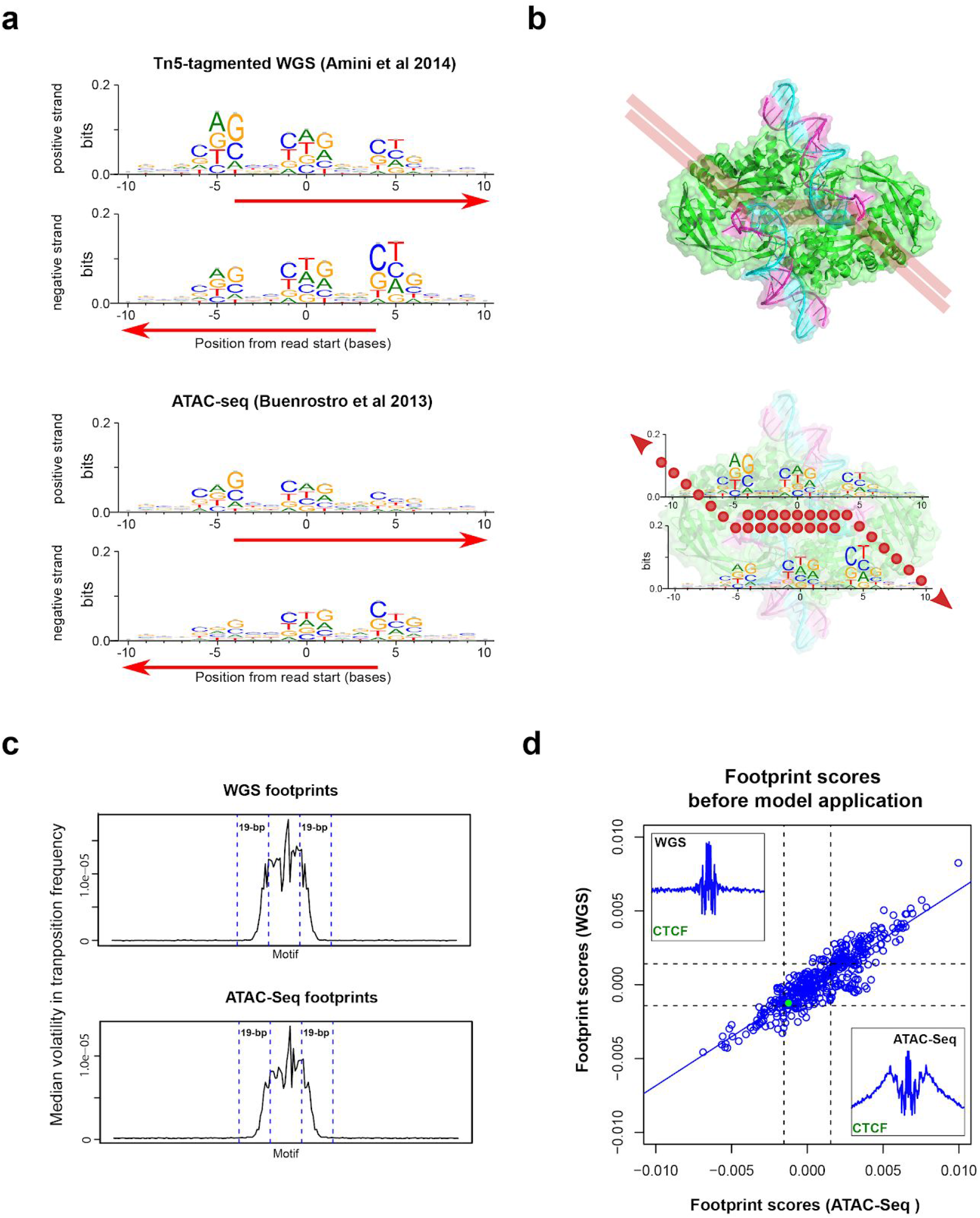
Sequence bias in Tn5 target sites confounds ATAC-Seq footprint estimates. **(a)** Observed 19-base palindromic sequence bias around read start sites of Tn5-tagmented WGS (top) and ATAC-seq (bottom). Only reads that mapped to genomic intervals defined as accessible regions in the ATAC-seq data were considered. Given that Tn5 introduces a 9 bp staggered cut, reads (red arrows) mapping to the positive and negative strand were shifted by +/− 4 bases respectively. **(b)** The crystal structure of the Tn5 homodimer complexed with the 19-bp mosaic end (ME) sequences is shown with the proposed path of the target DNA (red lines) on the transposome complex (top panel). The 3’ end of the ME DNA strands are transesterified to the target DNA with a 9-bp offset and the resulting sequence reads are shown in the bottom panel. Position-specific bias observed on the reads is depicted (lower panel) and likely arises due to interactions with neighboring residues on the transposome surface. **(c)** Transposition volatility is highly elevated within the motifs, and reduces to background levels 19-bases away from the motif. Transposition volatility at a position was defined as the square of the difference in the transposition frequency with the adjacent nucleotide position. All motifs were centered and transposition volatility was calculated for each motif. As motifs vary in length, the missing values were replaced by median volatility for the particular motif. The median volatility across all motifs is depicted here. **(d)** Transposition bias was tested in 471 non-redundant vertebrate motifs from the JASPAR CORE database. Footprints in motif instances observed in ATAC-seq exhibit a strong positive correlation (R^2^=0.80) with transposition patterns seen in instances found in WGS. Calculation of footprint scores is described in the Online Methods. Positive values indicate under-transposition (interpreted as footprints in ATAC-seq). Dotted lines indicate 1 standard deviation from the mean on each axis. (Plot insets) CTCF motif transposition observed in ATAC-seq (lower right corner) and WGS (upper left corner).

To assess if Tn5 transposition bias affected the determination of TF footprints in ATAC-seq data, we examine ATAC-seq and WGS libraries described above. Of note, these two libraries were prepared using the same hyperactive Tn5 transposase (Nextera DNA library preparation kit, Illumina). Only the TF binding motif matches that were present within the accessible intervals in the ATAC-seq data were considered. We observed that the average transposition frequency was highly volatile and varied greatly across the positions in the TF motif (Figure 4c). The stark contrast in the volatility of the transpositions within the TF motif and the flanking regions is a further indicator of Tn5 transposition bias, which is exaggerated upon aggregating a biased transposition signal across highly conserved motif matches in comparison to the flanking intervals that are not conserved. Considering the volatility in transposition within and adjacent to the motif matches, we smoothed the footprint traces by fitting a spline function with knots placed 19 bases away from the TF motif matches in line with the estimated 19-base span of the Tn5 transposition bias, and scored the footprints as the difference in transposition frequency at the knots placed in the flanking intervals and the center of the motif (Figure 4c,d). We calculated footprint scores for matches to all non-redundant vertebrate TF motifs present in the JASPAR CORE database and found a striking correlation (R^2^=0.801) between footprint scores in ATAC-Seq and WGS datasets (Figure 4d)^9^. As naked DNA is completely devoid of DNA-binding proteins, this indicates that the vast majority of footprints at TF binding motif matches in ATAC-Seq data are likely to be spurious and arise from the intrinsic bias of Tn5 transposition.

TF footprints in DNAse-Seq data have been previously shown to be confounded by positional bias arising from DNAse I cut site preferences in DNAse-seq data. While this bias can be corrected with the knowledge of preferred 6-mer cut sites used by DNAse I ^10^, such an approach is not scalable to the 19-mer bias observed at Tn5 transposition sites. The total number of possible 19-mer sequences far exceeds the number of bases within the genome by two orders of magnitude, so k-mer based representations of complex sequences biases would be both intractable and incomplete. However, we can train a Bayesian network to encode nucleotide dependencies at the sequences around ATAC-seq read start sites and thus learn the positional bias of Tn5 transposition. The parameters of the network must be restricted to the informative combinations of positions and nucleotides co-occurring at positions around the read start sites. Therefore we adopted Seqbias, an existing tool that was developed to correct positional bias in RNA-seq libraries using a Bayesian network model where the nodes are sequence positions surrounding read start sites and edges represent dependencies between the positions ^11^. The original article for Seqbias suggests that the underlying model could be applied to any short read data because it was built with very few assumptions and does not rely upon gene annotations. Here, we demonstrate that Seqbias can be used to correct Tn5 transposition bias in ATAC-Seq data, where Seqbias trains a Bayesian network model that can discriminate between the foreground sequences at the transposition sites from randomly sampled background sequences in their vicinity. Given that the Seqbias model is trained discriminatively, only parameters that are deemed informative in discriminating between foreground and background sequences are included in the model. Furthermore, Seqbias ignores duplicate reads and models only positional bias, in effect ignoring potential PCR biases that may be introduced during library amplification.

We used Seqbias to adjust the Tn5 transposition frequencies in both the WGS and ATAC-Seq datasets. The efficacy of Seqbias correction can be assessed by examining the nucleotide frequencies at the positions around the start sites of mapped reads (Figure 5a). The increased uniformity of nucleotide frequencies after correction is also evident from the shift in the Kullback-Leibler divergence calculated for each position around the transposition site with respect to a uniform background (Figure 5d). The transposition frequency across the bases of an individual instance of a motif at a genomic site may be too sparse to meaningfully estimate whether an individual site is occupied. Thus, we aggregated the Seqbias-adjusted transposition frequencies for each motif’s sites into a genome-wide trace, fitted a spline function to the trace, and computed a footprint score in order to obtain an estimate of the transcription factor’s genome-wide occupancy at instances of the motif.

**Figure 5:**
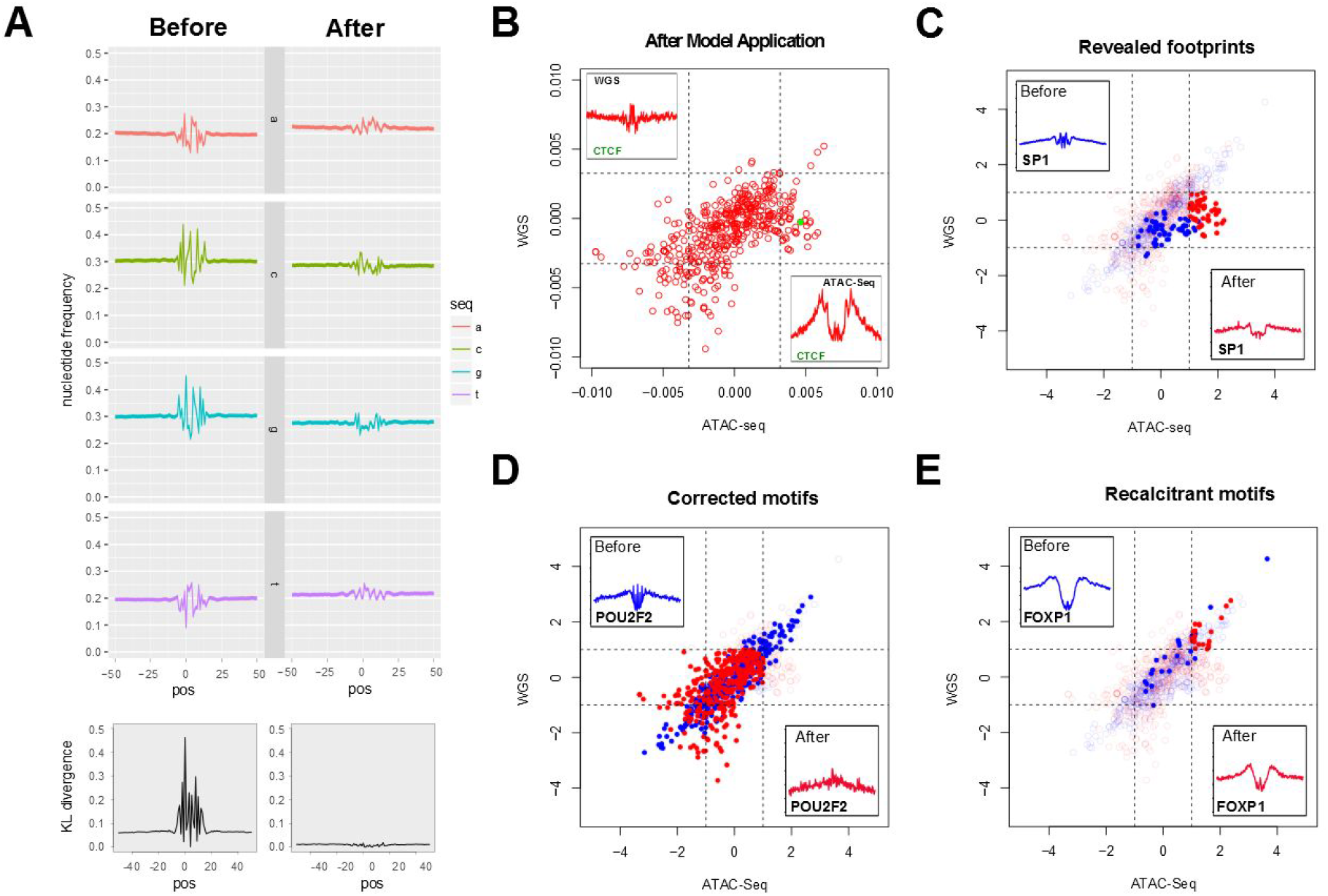
The Seqbias Bayesian network model yields corrected footprints. **(a)** (Top panel) Seqbias corrects nucleotide frequencies around read start sites to near-uniform levels. (Bottom panel) K-L divergence between nucleotide frequencies at positions around read start sites with respect to a uniform background distribution shown before (left) and after (right) Seqbias model application. **(b)** The correlation between the footprint scores observed in ATAC-seq and WGS data is reduced after Seqbias model application across the 471 JASPAR CORE motifs (R^2^=0.34). The number of matches for the motifs analyzed ranged from 88 (Rarg) to 20340 (NRF1) matches (median = 1080, IQR = 1498). Footprint scores were calculated as shown in Figure 3A. **(c)** 55 motifs (11.67%) are predicted to have genuine footprints in ATAC-seq data (e.g. SP1, shown in the inset). Highlighted points in (c), (d) and (e) indicate TF motif footprint Z-scores in ATAC-seq versus WGS before (blue) and red (after) model application. **(d)** 325 motifs (69.00%) are predicted not to be footprinted after Seqbias correction (e.g. POU2F2, shown in the inset). **(e)** 22 motifs (4.67%) show incomplete correction of transposition bias after model application (e.g. FOXP1, shown in the inset).

We observed that application of the model reduced the correlation between footprints observed in the ATAC-Seq and WGS datasets. Of the 471 motifs from the JASPAR CORE database tested, 352 motifs were determined as not footprinted in either the WGS or ATAC-Seq datasets (Figure 5c). 55 motifs were found to be footprinted only in the ATAC-Seq but not the WGS dataset. These TF motifs are therefore likely to be occupied in by cognate DNA binding proteins in native chromatin and represent genuine footprints (Figure 5d). 22 other motifs remain footprinted in both the ATAC-Seq and WGS datasets. These include what are likely to be spurious footprints as well as some motifs such as Foxp1 that are deeply footprinted in the WGS dataset but are recalcitrant to model correction (Figure 5e). While it is possible that some of these dually footprinted motifs may exhibit a true footprint in the ATAC-Seq data, the footprint estimates for this set of motifs are likely to be confounded by the background bias in transposition and additional methods will be necessary to assess if they are truly footprinted. We also examined how the footprints at the JASPAR CORE motif matches differed between GM12878 (B cells) and CD4^+^ T cells. TF motif matches in peaks accessible in either cell type were considered for this analysis and transposition events in the two cell types were counted across the same set of motif-containing intervals. Motifs that were footprinted in the WGS data were excluded. Among the footprints revealed by this analysis, 31 TF motifs were found to be footprinted in both cell types, 18 motifs were footprinted exclusively in GM12878 cells and 9 were footprinted in CD4+ T cells alone **(Figure S2)**. A complete list of TF footprints calculated using this approach is included in the Supplementary Material **(Table S1)**.

Our approach to adjust Tn5 transposition bias and generate a global estimate of transcription factor binding site occupancy is easily generalizable and can be used to correct footprinting data from other types of chromatin accessibility experiments such as DNAse-Seq. We also note that estimates of ATAC-seq peak heights may be affected by transposition bias but given that ATAC-seq peaks are up to two orders of magnitude longer than typical TF motifs, the sequence bias is likely to be averaged out within individual peaks. We believe that functional changes in transcription factor occupancy can be best discerned from differences in ATAC-seq footprints observed within the same set of genomic intervals upon biological perturbations. Identification of TF binding sites in accessible regions of ATAC-Seq data using *de novo* footprinting is not possible without due consideration of Tn5 transposition bias. Indeed, a high proportion of ATAC-seq footprints at TF binding motif matches spuriously arise from intrinsic Tn5 transposition bias rather than from specific DNA-protein interactions.

## METHODS

### Alignment and peak calling

ATAC-seq data (SRR891268) and WGS data (SRR1554094) used in this study were downloaded from NCBI. The ATAC-Seq library was generated using a human B lymphoblastoid cell line (GM12878) and CD4^+^ T cells^1^. The WGS library was prepared using human genomic DNA ^12^. Reads were aligned using Bowtie2 (v2.2.4) to the hg38 reference genome using default parameters, and a MAPQ cutoff of 1 was used. Accessible regions in the ATAC-seq data were determined using MACS2 (v2.1.1 in the “narrow” mode using default parameters). Examination of motif instances and transposition patterns in the ATAC-seq and WGS data was restricted to these accessible genomic intervals.

### Identification of motif instances

Nonredundant vertebrate TF motifs were downloaded from the JASPAR CORE database as position frequency matrices (PFMs), converted to the MEME format, and FIMO was used to determine instances of the motifs in the accessible genomic intervals (motif matches with p-value < 1e-04 were considered) ^9,13^. 100 bp on either side of motif matches were considered for footprinting analysis. Motifs with inadequate coverage were excluded from further analysis.

### Estimating Tn5 positional bias

20-base sequences flanking all read start sites mapping within the accessible peak intervals were obtained. Duplicate reads were ignored and reads mapping to the positive and negative strands were analyzed separately. Read starts mapping to the positive and negative strand were shifted by +/− 4 bases respectively to account for 9-base staggered cut introduced by Tn5. WebLogo was used to visualize the consensus sequence surrounding the read start sites ^14^

### Testing CENTIPEDE on simulated reads

Briefly, CENTIPEDE assesses the likelihood of a putative motif-containing genomic interval exhibiting a particular transposition pattern, given a prior probability of it being bound or not, using a hierarchical mixture model that considers the bound and unbound states of the motif^3^. The read coverage for motif-containing intervals is modeled as a negative binomial distribution, while the positional distribution of reads within individual motif-containing intervals is modeled as a multinomial distribution. The product of this likelihood function and the prior probability of a motif being bound yields a posterior probability, which is used to predict whether a particular instance of a motif is bound or not.

We generated a matrix of insertion site counts for Smad3 and Tbx5, two transcription factor motifs that do not exhibit footprints (even after Seqbias correction) in the GM12878 ATAC-seq dataset. We ran CENTIPEDE using default runtime parameters without specifying PWM match scores or motif conservation scores as we were interested in direct evidence of TF occupancy based upon steric exclusion of transposition events (Figure 1). To test CENTIPEDE on cases that may yield spurious predictions of footprints, we simulated cut site counts across a set of intervals bearing non-footprinted motifs using R. The total number of cut sites per interval (coverage) was sampled from a uniform distribution ranging from zero to a maximum of 400. The intervals were divided into two sets, one with a uniform distribution of cut-sites and a second with a non-uniform (normal) distribution of cut sites centered on the motif, and the cut-sites were distributed across each interval accordingly (Figure 2). Two sets of data were simulated: (i) with no difference and (ii) with a 2-fold difference in the mean coverage between the intervals bearing a uniform and a non-uniform positional distribution, and CENTIPEDE was run as described above.

### Calculation of footprint scores

For footprint visualization and downstream processing, position-adjusted positive- and negative-stranded transposition tracks were added together. To create a symmetric footprint trace, the transposition frequency track across the motif containing intervals was added to the reverse of itself and divided by 2. Volatility at each motif position was defined as the square of the differences in transposition frequencies of successive nucleotide positions (a position offset of 1). All motifs were centered and transposition volatility was calculated across each motif. As motifs vary in length, the missing values for shorter motifs were replaced by the median volatility for the particular motif. The median volatility across all motifs is depicted in Figure 4C. We found that a spline model fit using the midpoint of the motif, the ends of the motif match, and positions 19 bases away from the motif as knots was robust to extremes in transposition frequencies within the motif interval (Figure 3A). The choice of 19 bases reflects the span of the estimated Tn5 sequence bias surrounding a read start site. Further, the volatility declined to background levels over this distance (Figure 4C). The footprint score was defined as the difference between the modeled transposition frequencies at the knots corresponding to the 19-base flank and the midpoint of the motif. Footprint scores were not correlated with transposition frequencies at the spline knot positions placed at the 19-base flanks, implying that the footprint scores were not associated with coverage (Figure 3B). In this study, motifs with a footprint Z-score > 1 were considered to be footprinted.

### Seqbias

For each motif, a BED file containing motif matches expanded by 100 bp on either side was imported into Seqbias, along with the BAM alignment files from the ATAC-seq and WGS datasets. Transposition events in these intervals, were estimated by counting the read start sites using the ‘binary=TRUE’ mode, where a position has a count of 0 if no read maps to it and 1 if at least one does; this counting scheme produces a more robust footprint estimate, devoid of amplification bias. The Seqbias model used for correcting ATAC-seq read counts was trained on the ATAC-seq data, with sequences 20 bases on either site of each read start site considered for the model. For adjusting the WGS read counts, a separate Seqbias model was trained on reads mapping to chromosome 22, comprising ~3.6 million mapped reads. Since the performance of Seqbias appears to level off at ~10^5^ reads, the model trained on chromosome 22 can be expected to perform comparably to a model built using all reads in the dataset ^11^. Given that the reads mapping to the positive and negative strands exhibit an identical bias, all reads can be corrected with the same model. We also compared the use of a common Seqbias model trained on read start sites exclusive to chromosome 22 with motif-specific models trained on WGS reads limited to the intervals containing matches for each motif across the entire genome, but did not obtain in any further improvement in performance (data not shown). The Seqbias models used for bias correction are available as YAML files (Supplementary Data). The overall approach for Seqbias-corrected TF footprints is summarized in the flowchart below (Figure S3).

## Additional Information

### Author contributions

VM, VV and SP conceived the project. VM and VV and carried out the analysis and wrote the manuscript. SP reviewed the text.

### Competing financial interests

The authors have no potential conflicts of interest to declare.

