## Supplemental Materials for "A Bayesian approach for correcting Tn5 transposition bias in ATAC-seq footprinting"

### SUPPLEMENTARY FIGURES

**Figure S1: Tn5 bias is retained at different distances from the summit of accessibility peaks**

**Figure S2: Motifs exclusively footprinted in CD4+ T naive cells or GM12878 B cell line are revealed by application of the approach**

**Figure S3: A schematic overview of the approach used to correct footprint estimates.**

### SUPPLEMENTARY TABLE

**Table S1: JASPAR CORE transcription factor motifs for which Tn5 bias was corrected and footprint scores were calculated in Figure S2.**

**Figure S1: Tn5 bias is retained at different distances from the summit of accessibility peaks.** The motif is calculated for ATAC-seq read start sites within (left) 50 bases and (right) 100 bases of the peak summits. The controls shown were calculated after *randomly* offsetting insertion sites by up to 10 bp.

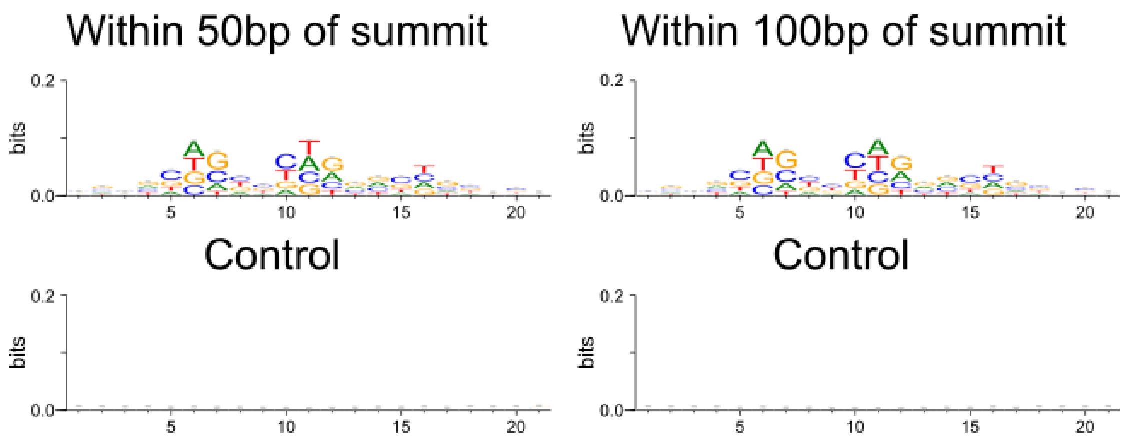

**Figure S2: Motifs exclusively footprinted in CD4+ T naive cells or GGM1287 B cell line are revealed by application of the Seqbias model.** Footprints are plotted as Z-scores. Motifs that are not footprinted in the WGS data are considered for further analysis. Among these, footprints that are footprinted (Z-score > 1) in T and/or B cells are identified (marked in color). The Seqbias-corrected footprint scores are listed in the accompanying table.

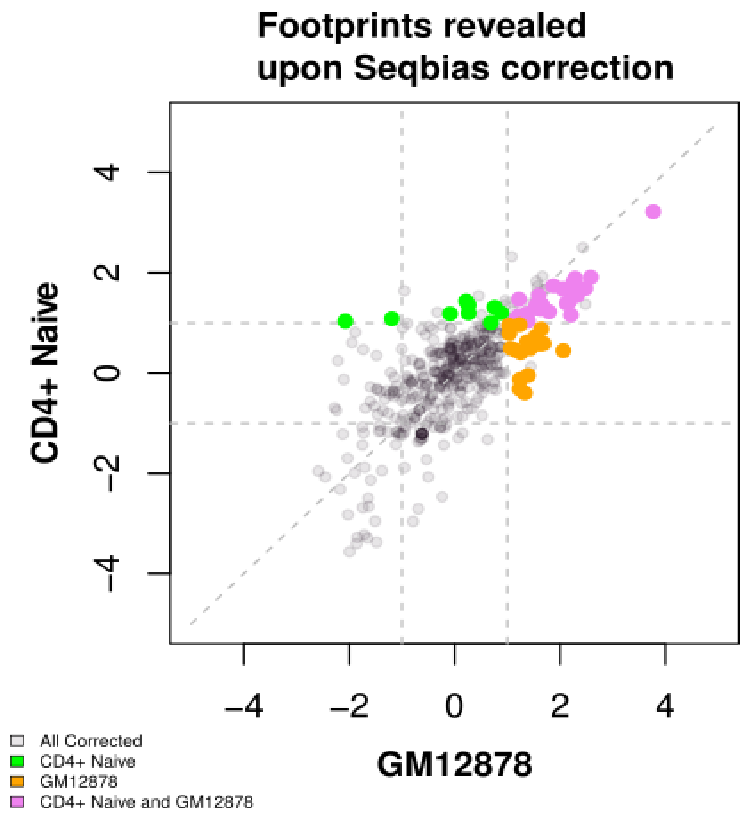

**Figure S3: A schematic overview of the use of Seqbias to correct footprint estimates.**

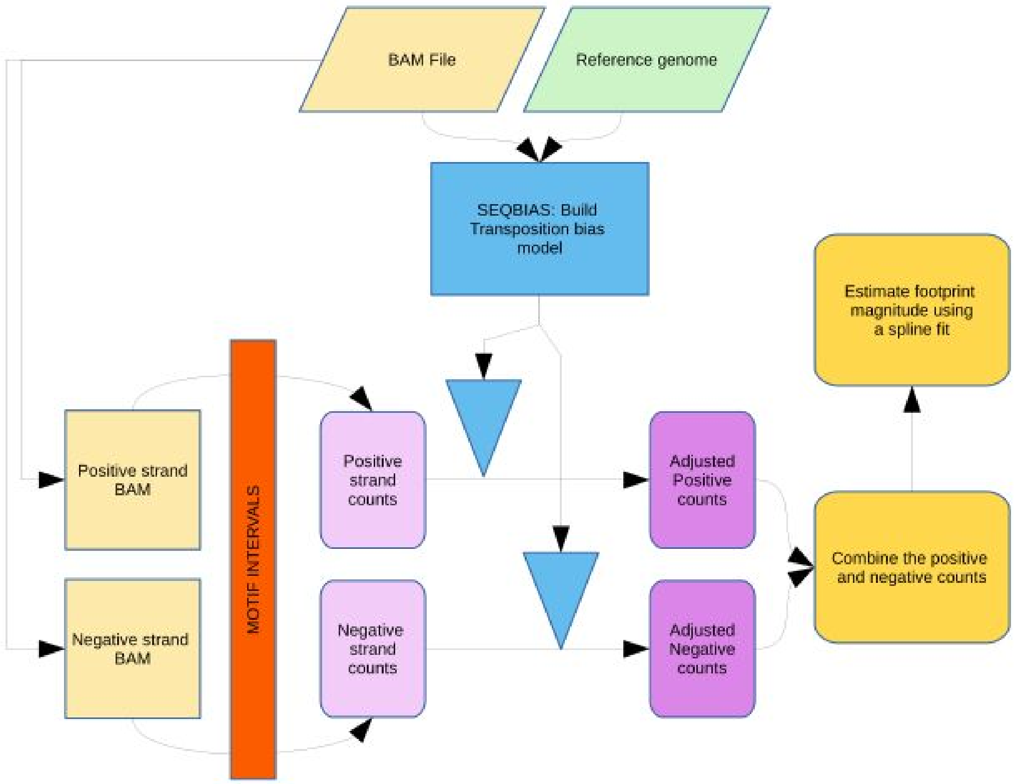

**Table S1: JASPAR CORE transcription factor motifs for which Tn5 bias was corrected and footprint scores were calculated in Figure S2.**

| Footprinted Motifs | Footprint Score (Seqbias-corrected) | |
|---|---|---|
| In both T and B cells | B cells (GGM1287) | CD4+ naive T cells |
| MA0079.3_SP1 | 1.558058 | 1.367833 |
| MA0093.2_USF1 | 2.124897 | 1.392325 |
| MA0139.1_CTCF | 3.770354 | 3.218626 |
| MA0162.2_EGR1 | 1.396466 | 1.041543 |
| MA0464.2_BHLHE40 | 2.052229 | 1.676906 |
| MA0472.2_EGR2 | 1.368315 | 1.110769 |
| MA0496.1_MAFK | 1.220874 | 1.473267 |
| MA0516.1_SP2 | 1.209592 | 1.13577 |
| MA0526.1_USF2 | 1.79378 | 1.221952 |
| MA0603.1_Arntl | 2.289165 | 1.610728 |
| MA0606.1_NFAT5 | 1.630446 | 1.276699 |
| MA0620.1_Mitf | 2.487472 | 1.692165 |
| MA0625.1_NFATC3 | 2.209368 | 1.157743 |
| MA0636.1_BHLHE41 | 2.583748 | 1.912693 |
| MA0657.1_KLF13 | 1.202809 | 1.020732 |
| MA0663.1_MLX | 2.307269 | 1.563993 |
| MA0664.1_MLXIPL | 2.317999 | 1.546921 |
| MA0685.1_SP4 | 1.872213 | 1.737802 |
| MA0692.1_TFEB | 2.189169 | 1.441127 |
| MA0728.1_Nr2f6(var.2) | 1.091963 | 1.04277 |
| MA0732.1_EGR3 | 1.369187 | 1.185492 |
| MA0733.1_EGR4 | 1.348732 | 1.083555 |
| MA0740.1_KLF14 | 1.652991 | 1.367311 |
| MA0741.1_KLF16 | 1.616933 | 1.38533 |
| MA0742.1_Klf12 | 1.599042 | 1.550243 |
| MA0746.1_SP3 | 1.640651 | 1.33894 |
| MA0747.1_SP8 | 1.543505 | 1.293153 |
| MA0828.1_SREBF2(var.2) | 2.25322 | 1.820174 |
| MA0829.1_Srebf1(var.2) | 2.286336 | 1.891391 |
| MA0831.1_TFE3 | 2.267188 | 1.615395 |
| MA0871.1_TFEC | 2.398235 | 1.669322 |
| Only in T cells | B cells (GGM1287) | CD4+ naive T cells |
| MA0117.2_Mafb | 0.26616604 | 1.195393 |
| MA0493.1_Klf1 | 0.68800492 | 1.006094 |
| MA0628.1_POU6F1 | 0.21944715 | 1.434456 |
| MA0681.1_Phox2b | -2.07400736 | 1.038082 |
| MA0715.1_PROP1 | -1.19113224 | 1.085779 |
| MA0769.1_Tcf7 | -0.08628684 | 1.18242 |
| MA0836.1_CEBPD | 0.88310455 | 1.199816 |
| MA0837.1_CEBPE | 0.75964096 | 1.311789 |
| MA0899.1_HOXA10 | 0.26483007 | 1.358774 |
| Only in B cells | B cells (GGM1287) | CD4+ naive T cells |
| MA0018.2_CREB1 | 1.379033 | 0.62273849 |
| MA0041.1_Foxd3 | 1.234481 | -0.31706374 |
| MA0046.2_HNF1A | 1.241205 | -0.12519975 |
| MA0101.1_REL | 1.154856 | 0.45639155 |
| MA0104.3_Mycn | 1.03714 | 0.79429156 |
| MA0105.4_NFKB1 | 1.550584 | 0.74003717 |
| MA0107.1_RELA | 1.438197 | 0.48826221 |
| MA0154.3_EBF1 | 1.33239 | -0.39665893 |
| MA0469.2_E2F3 | 1.031903 | 0.95293828 |
| MA0497.1_MEF2C | 1.396144 | -0.04612146 |
| MA0609.1_Crem | 1.251251 | 0.40548854 |
| MA0624.1_NFATC1 | 2.061727 | 0.44611698 |
| MA0656.1_JDP2(var.2) | 1.504577 | 0.58044736 |
| MA0753.1_ZNF740 | 1.231007 | 0.96185538 |
| MA0778.1_NFKB2 | 1.642954 | 0.87677537 |
| MA0834.1_ATF7 | 1.685167 | 0.58910827 |
| MA0835.1_BATF3 | 1.069457 | 0.49560462 |
| MA0840.1_Creb5 | 1.618581 | 0.58130703 |
